# The STORE.2 model of the T cell proliferative phase considering c-Myc

**DOI:** 10.1101/2025.06.05.658086

**Authors:** David A. Christian, Taran P.S. Bhartt, Zachary Lanzar, Seyedeh Fatemeh Seyyedizadeh, Christopher A. Hunter, Thomas A. Adams

**Affiliations:** University of Pennsylvania, Department of Pathobiology, Philadelphia, Pennsylvania, USA; McMaster University, Department of Chemical Engineering, Hamilton, Ontario, Canada; Norwegian University of Science and Technology, Department of Energy and Process Engineering, Trondheim, Norway

**Keywords:** T cell, Modelling, c-Myc, proliferative phase, vaccine

## Abstract

The generation of antigen-specific CD8^+^ T cell responses is dictated by the affinity of the cognate antigen, the stimulatory capacity of antigen presenting cells (APCs), and the metabolic pathways required for rapid cell proliferation. The complexity of these pathways is a significant challenge in designing vaccines against diseases that require a protective CD8^+^ T cell response. To understand the mechanisms underlying CD8^+^ T cell responses, the STORE.2 model was developed to simulate the early events of T cell priming and expansion at the site of priming. STORE.2 is a mathematical, stochastic, and agent-based model based on first-principles that tracks every individual CD8^+^ T cell and APC. It allows for the simulation of different antigen affinity (Signal 1) as well as levels of costimulation (Signal 2) and inflammatory cytokines (Signal 3) provided by APCs. The impact of Signals 1-3 is translated to T cell responses via the transcription factor c-Myc, which supports T cell proliferation. Enhanced glycolysis during T cell activation results in a change in the metabolic environment that includes an increase in extracellular lactate concentration. STORE.2 models the role of lactate in the inhibition of T cell metabolism and proliferation as a negative feedback mechanism on c-Myc production. STORE.2 accurately recapitulated the CD8^+^ T cell response during *in vitro* T cell priming assays that control for Signals 1-3 as well as lactate concentration. Finally, application of STORE.2 to an *in vivo* response to immunization demonstrated that the model accurately simulates CD8^+^ T cell activation at the site of priming.

## INTRODUCTION

CD8^+^ T cells provide a critical role in protection against a variety of intracellular pathogens such as HIV (*1*), tuberculosis (*2*), malaria (*3*), *Toxoplasma gondii* (*4*), and COVID-19 (*5, 6*) as well as in anti-tumor immunity (*7*). This protective capacity has motivated efforts to develop vaccines that can elicit robust CD8^+^ T cell responses (*8*– *10*). However, the cellular signaling cascades required to drive CD8^+^ T cell clonal expansion and memory T cell development are exceedingly complex and have limited progress in the generation of vaccine formulations that reliably generate protective CD8^+^ T cell memory (*9*). Computational models can provide clarity by rapidly simulating these dynamics, tracking millions of cell-cell interactions and cellular processes over time in ways that are difficult experimentally (*11*–*13*).

Computational models have been used to examine key aspects of CD8^+^ T cell responses, including control of proliferation, cell death, clonal expansion, and T cell differentiation (*11, 14*). Early work was able to capture the dynamics of T cell activation and expansion *in vitro* by using population-based models (*15*–*18*). For example, the Cyton model determined the division or death times for lymphocytes using separate probability distributions (*19*).

Importantly, this model introduced the concept of Division Destiny (DD) to account for the variation in number of divisions lymphocytes will undergo before returning to quiescence or dying. The Cyton model then helped establish that the initial stimulation level of lymphocytes determined the DD of the population via a heritable factor (*15, 20*–*25*). This idea motivated the development of the agent-based Cyton2 model to track these heritable factors in specific T cell clones (*26*). One such heritable factor was found to be the transcription factor c-Myc, the expression of which was shown to scale with the level of stimulus of lymphocytes and had a required threshold to allow for continued cell division (*24*).

c-Myc is upregulated immediately upon T cell receptor (TCR) stimulation and is required for the subsequent metabolic reprogramming of T cells that allows for rapid T cell proliferation and differentiation (*27, 28*). In addition to being required for the upregulation of glucose and glutamine transporters that aid in glycolysis and glutamine catabolism (*27*), c-Myc regulates the expression of amino acid (AA) transporters (*29*) needed for protein synthesis and T cell proliferation (*30, 31*). c-Myc also serves critical roles in cell cycle phase progression in other cell types by controlling the transitions between different cell cycle phases (*32*) as well by regulating other proteins that control cell cycle (*33*). Cellular levels of c-Myc are maintained by first inducing c-Myc mRNA from stimulatory factors such as TCR engagement (Signal 1) and signaling from costimulatory molecules (Signal 2) and cytokines (Signal 3) (*24, 28, 34*). c-Myc protein is then controlled by post-translational modifications that control the rate of c-Myc protein degradation (*34, 35*) as well as the ability to take up AA to translate c-Myc protein (*31, 34*). In addition to stimulating c-Myc mRNA, cytokines have been shown to enhance c-Myc levels by enhancing uptake of AA and protein synthesis (*34*) and blocking c-Myc protein degradation through O-GlcNAcylation (*36, 37*). The multifaceted role that c-Myc plays in T cell metabolism, protein synthesis, and potentially cell cycle control motivate modeling its expression level and its impact on the T cell response.

In prior work, we developed the STochastic Omentum REsponse (STORE.1) model, an agent-based stochastic finite-state automata used to simulate APC priming of antigen-specific CD8^+^ T cell clonal expansion *in vivo* (*38*). The model replicated experiments in which mice first received transgenic OT-I CD8^+^ T cells specific for ovalbumin (OVA) followed by intraperitoneal (i.p.) vaccination with an OVA-secreting vaccine strain of *Toxoplasma gondii* (CPS-OVA). The model considers how long it takes for each naive OT-I T cell to find an antigen presenting cell (APC) presenting its cognate antigen after i.p. vaccination and then tracks the subsequent T cell divisions over time. A follow-up study considered a similar experiment in which OT-I T cell activation in the spleen was tracked in response to an adjuvanted subunit vaccine administered intravenously, and demonstrated that the model could explain experimental data when applied to the different tissues and vaccine formulations (*39*).

To determine the time required for T cell divisions, the STORE.1 model used probability distributions based on experimental observations in the literature about the length of the cell cycle at different points in the activation process. However, these distributions contained no underlying mechanism as to why the cell cycle duration changes with division number. To address this limitation, a new fundamental cellular mechanism was added to track the c-Myc level of individual T cells from priming and activation by APCs through the proliferative phase. After the initial induction of c-Myc by TCR engagement, its expression level is amplified by APCs via Signals 2 and 3. Subsequently, AA uptake that is increased by c-Myc expression serves to increase the rate of c-Myc production. In contrast, c-Myc induced glycolysis generates lactate that impairs metabolic pathways required to produce more c-Myc (*40*–*44*). As c-Myc levels control the metabolic and AA uptake pathways required for rapid cell division, the progression of every T cell through the cell cycle was modeled as a function of c-Myc expression level rather than by the probability distributions used in STORE.1. These dynamic feedback loops provide a mechanistic explanation of how c-Myc levels degrade with increasing divisions and result in longer cell cycle times until c-Myc levels decrease to a level that results in a cessation of division. In summary, the STORE.2 model presented here is an agent-based model (ABM) of T cell activation that accurately simulates the impact of T cell extrinsic pathways on the generation of CD8^+^ T cell responses to vaccination.

## RESULTS

### STORE.2 Model

Like the STORE.1 model (Fig. 1A), the STORE.2 model (Fig. 1B) is a stochastic, ABM of the early priming and expansion phase of the CD8^+^ T cell response in the omentum after intraperitoneal immunization with the attenuated *ΔcpsII* strain of the parasite *Toxoplasma gondii* (*38*). The steps of the process of T cell priming were parameterized such that naive OT-I T cells (cell type *A*) enter the omentum at a constant rate over 24 hr. In STORE.2, these naive cells can now also randomly leave the system after 24 hr at a modeled rate of *r*_*A,leave*_:

**Figure 1.**
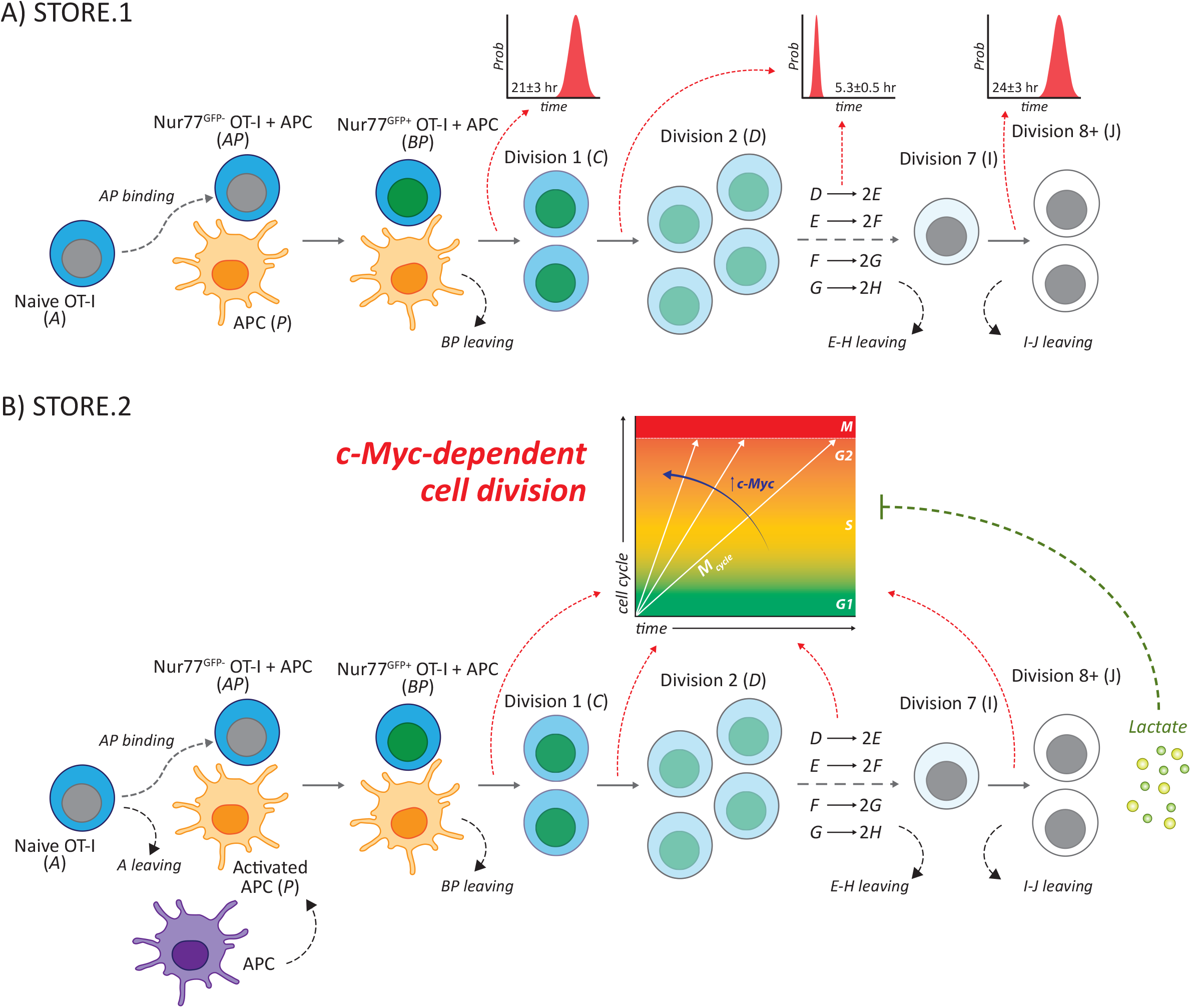
STORE.1 and STORE.2 models. (**A**) STORE.1 models the arrival time and priming of naive OT-I T cells (*A*) by any APC (*P*) in the omentum. Naive T cells bind to an APC (*AP* cell) and become activated (*BP* cell) based on the expression of the Nur77^GFP^ TCR reporter. Activated T cells then divide into two daughter cells (*C* cells) at a rate determined by a probability distribution function (PDF). Subsequent divisions (*C*-*H* cells and *I*-*J* cells) also occur at rates determined by two different PDFs. Cell types *BP, E*-*H*, and *I*-*J* can all leave the system at 3 different rates that were determined by the parameter estimation algorithm. (**B**) STORE.2 modifies STORE.1 by first accounting for APC activation times and then allowing only activated APCs to bind and prime naive T cells. The PDFs used for cell division rates are replaced by a c-Myc-dependent cell division function that is applied to all cell divisions. STORE.2 tracks the glycolytic byproduct lactate and its ability to impair c-Myc expression and slow cell division.

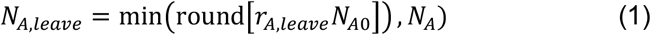

In STORE.1, APCs (cell type *P*) entered the system at a constant rate and could interact with and activate naive T cells. The STORE.2 model now accounts for the interaction between APCs and the parasite. This interaction is required for the APC to present the cognate antigen to T cells (Signal 1) and to become activated in order to express costimulatory molecules (Signal 2) and pro-inflammatory cytokines (Signal 3). The time required for APCs to activate (*t*_*act*_) and be designated as a *P* cell that is available for binding to naive T cells is based on a normal probability distribution function (PDF):

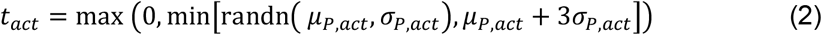

where *μ*_*P,act*_ and *σ*_*P,act*_ are the mean and standard deviation activation times and the upper bound of *t*_*act*_ is three standard deviations longer than the mean.

The binding of each activated APC to a naive T cell is determined at each timestep based on the probability:

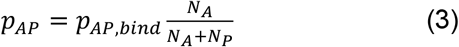

where *p*_*AP,bind*_ is a constant parameter, and *N*_*A*_ and *N*_*P*_ are the numbers of *A* and *P* cells, respectively. When *A* and *P* cells initially bind, they transition to *AP* cells (APCs bound to Nur77^GFP-^ OT-I T cells). The signaling of the cognate antigen presented by the APC via major histocompatibility complex I (MHC-I) to the T cell receptor (TCR) on the naive T cell is then tracked by the expression of the Nur77^GFP^ reporter. The Nur77^GFP+^ OT-I T cell bound to an APC is designated as cell type *BP*. The time of this *AP* to *BP* transition is based on a normal PDF (*μ* = 30 min, *σ* = 10 min) that approximates the time for GFP to be expressed at detectable levels.

The process of T cell priming on an APC has been shown to include three phases where the longest phase is characterized by APC:T cell contacts that last approximately 24 hours before T cells become migratory and begin dividing rapidly. This phase was modeled in STORE.1 using random number generation against a normal PDF (*μ* = 21 hr, *σ* = 3 hr) that resulted in:

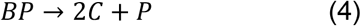

Subsequent cell divisions were similarly modeled using normal PDFs with means of 5.3 hours for cells in divisions 1 through 6 (cell types *C* through *H*) and 24 hours for divisions 7 and 8+ (cell types *I* and *J*). In STORE.2, cell division is determined by progress through cell cycle that is tracked in the model. The speed that cells go through cell cycle is controlled by the transcription factor c-Myc that is first induced during T cell activation and then has its concentration in each T cell determined by feedback loops.

To account for cells leaving the system by either cell death or migrating out of the omentum in STORE.1, each *BP* and *E* through *J* cell types are checked at each time point with a random number generator versus a normal PDF. The model uses parameters *p*_*BP,leave*_, *p*_*EH,leave*_, and *p*_*IJ,leave*_ to describe the percentage chance that cell types *BP, E* through *H*, and *I* through *J* leave during that timestep, respectively. In STORE.2, the leaving rate of *BP* cells (*p*_*BP,leave*_) is simplified to a fixed rate such that the number that leave at each timestep is determined by *round*(*p*_*BP,leave*_ *N*_*BP*_) where *N*_*BP*_ is the number of *BP* cells at that timestep.

### Signal 1 drives initial c-Myc synthesis

The activation of T cells by Signal 1 (Fig. 2A) has been shown to induce rapid expression of c-Myc protein dependent upon the TCR affinity to the peptide:MHC-I complex (*28*). To measure this induction experimentally, OT-I/Myc^GFP/GFP^ mice were generated by crossbreeding OT-I mice with Myc^GFP/GFP^ reporter mice. OT-I T cells were isolated and stimulated *in vitro* using either the high affinity SIINFEKL peptide (N4) or the low affinity SIIVFEKL peptide (V4) (*45, 46*). The induction of c-Myc protein expression by Signal 1 was measured at 24 hr after stimulation by Myc^GFP^ fluorescence intensity and showed that TCR stimulation with the V4 peptide:MHC-I induced 39±7% as much c-Myc as the N4 peptide (Fig. 2B, C). To capture this *in vitro* priming in the model, the introduction rate of *A* and *P* cells was decreased to 1 hr to correspond to a faster arrival time than modeled for *in vivo* immunizations, and *p*_*BP,leave*_ was set to zero to account for self-priming *in vitro* as *r*_*A,leave*_ was nonzero. In addition, the *in vitro N*_*A*0_/*N*_*P*0_ ratio and the APC activation time parameters *μ*_*P,act*_ and *σ*_*P,act*_ were estimated by the parameter estimation algorithm (*Methods*, Table 1). Upon TCR stimulation and formation of an AP cell, c-Myc expression by T cells was modeled as a step change increase in the number of c-Myc proteins contained in the T cell. This step change was randomly chosen from a normal distribution of c-Myc protein number found by parameter estimation for both peptides (N4: *μ*_*cMYC*0_ = 3500, *σ*_*cMYC*0_ = 2500; V4: *μ*_*cMYC*0_ = 1500, *σ*_*cMYC*0_ = 1500). The average c-Myc protein number in T cells was then measured at 24 hr and showed that V4 peptide stimulation resulted in 61±3% as much c-Myc protein as stimulation with N4 peptide (Fig. 2D). To further account for the decrease in V4 peptide affinity and subsequent decrease in activation efficiency, the ratio of naive T cells and APCs input into the system (*N*_*A*0_/*N*_*P*0_) was increased for the V4 peptide. These data demonstrate that the variable step change in the model for c-Myc protein expression based on peptide affinity captured the difference in c-Myc levels observed experimentally at early time points.

**Table 1.**
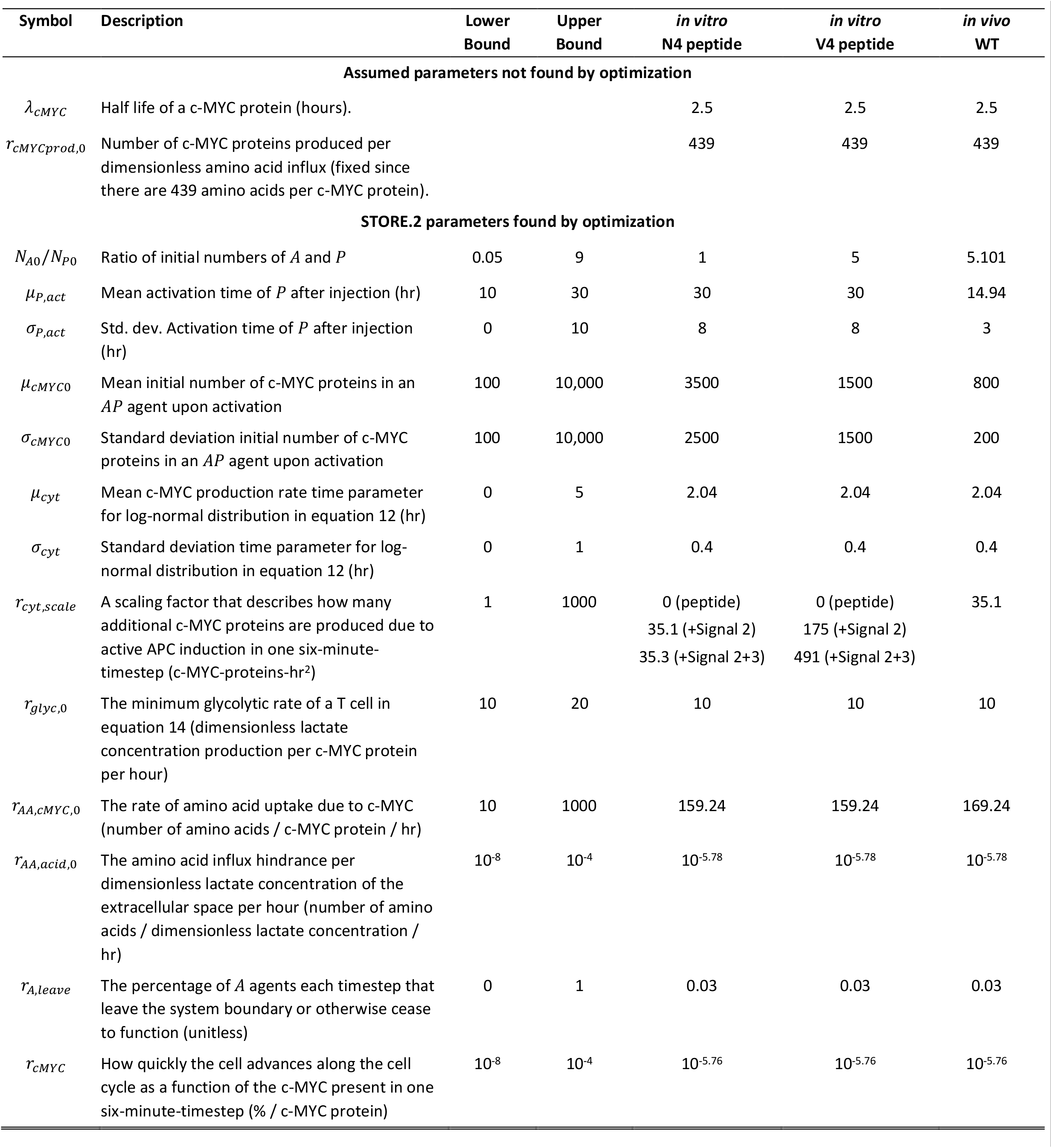
Final recommended STORE.2 model parameters and bounds used in the parameter estimation optimization problem (see *Methods*).

| Symbol | Description | Lower Bound | Upper Bound | *in vitro*<br>N4 peptide | *in vitro*<br>V4 peptide | *in vivo*<br>WT |
| --- | --- | --- | --- | --- | --- | --- |
| **Assumed parameters not found by optimization** |  |  |  |  |  |  |
| $\lambda_{cMYC}$ | Half life of a c-MYC protein (hours). | | | 2.5 | 2.5 | 2.5 |
| $r_{cMYCprod,0}$ | Number of c-MYC proteins produced per dimensionless amino acid influx (fixed since there are 439 amino acids per c-MYC protein). | | | 439 | 439 | 439 |
| **STORE.2 parameters found by optimization** |  |  |  |  |  |  |
| $N_{A0}/N_{P0}$ | Ratio of initial numbers of *A* and *P* | 0.05 | 9 | 1 | 5 | 5.101 |
| $\mu_{P,act}$ | Mean activation time of *P* after injection (hr) | 10 | 30 | 30 | 30 | 14.94 |
| $\sigma_{P,act}$ | Std. dev. Activation time of *P* after injection (hr) | 0 | 10 | 8 | 8 | 3 |
| $\mu_{cMYC0}$ | Mean initial number of c-MYC proteins in an *AP* agent upon activation | 100 | 10,000 | 3500 | 1500 | 800 |
| $\sigma_{cMYC0}$ | Standard deviation initial number of c-MYC proteins in an *AP* agent upon activation | 100 | 10,000 | 2500 | 1500 | 200 |
| $\mu_{cyt}$ | Mean c-MYC production rate time parameter for log-normal distribution in equation 12 (hr) | 0 | 5 | 2.04 | 2.04 | 2.04 |
| $\sigma_{cyt}$ | Standard deviation time parameter for log-normal distribution in equation 12 (hr) | 0 | 1 | 0.4 | 0.4 | 0.4 |
| $r_{cyt,scale}$ | A scaling factor that describes how many additional c-MYC proteins are produced due to active APC induction in one six-minute-timestep (c-MYC-proteins-hr<sup>2</sup>) | 1 | 1000 | 0 (peptide)<br>35.1 (+Signal 2)<br>35.3 (+Signal 2+3) | 0 (peptide)<br>175 (+Signal 2)<br>491 (+Signal 2+3) | 35.1 |
| $r_{glyc,0}$ | The minimum glycolytic rate of a T cell in equation 14 (dimensionless lactate concentration production per c-MYC protein per hour) | 10 | 20 | 10 | 10 | 10 |
| $r_{AA,cMYC,0}$ | The rate of amino acid uptake due to c-MYC (number of amino acids / c-MYC protein / hr) | 10 | 1000 | 159.24 | 159.24 | 169.24 |
| $r_{AA,acid,0}$ | The amino acid influx hindrance per dimensionless lactate concentration of the extracellular space per hour (number of amino acids / dimensionless lactate concentration / hr) | $10^{-8}$ | $10^{-4}$ | $10^{-5.78}$ | $10^{-5.78}$ | $10^{-5.78}$ |
| $r_{A,leave}$ | The percentage of *A* agents each timestep that leave the system boundary or otherwise cease to function (unitless) | 0 | 1 | 0.03 | 0.03 | 0.03 |
| $r_{cMYC}$ | How quickly the cell advances along the cell cycle as a function of the c-MYC present in one six-minute-timestep (% / c-MYC protein) | $10^{-8}$ | $10^{-4}$ | $10^{-5.76}$ | $10^{-5.76}$ | $10^{-5.76}$ |

**Figure 2.**
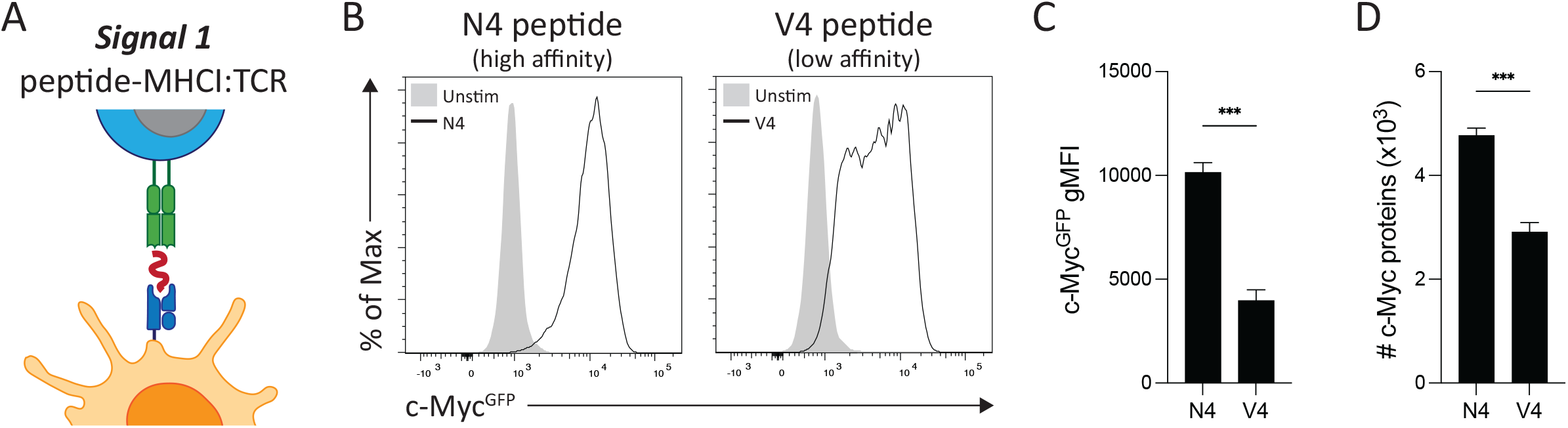
Impact of TCR signal strength on c-Myc expression. (**A**) Schematic of the Signal 1 peptide:MHCI antigen presentation to a CD8^+^ T cell. (**B**) Flow cytometry analysis of c-Myc^GFP^ fluorescence intensity in OT-I/Myc^GFP/GFP^ T cells stimulated with 20 ng of either N4 or V4 peptide *in vitro* for 24 hr versus unstimulated controls. (**C**) Geometric mean fluorescence intensity (gMFI) of c-Myc^GFP^ expression measured for N4 and V4 peptide stimulation at 24 hr. (**D**) STORE.2 simulation results for c-Myc protein number per OT-I T cell stimulated with N4 or V4 peptide for 24 hr. Data are representative plots from (B and C) three experiments or (D) three simulation runs. Bar graphs indicate means ± SD. Data analyzed by unpaired *t* test. \*\*\**P* < 0.001.

### c-Myc expression control by APC activation and feedback loops

The expression level of c-Myc after Signal 1 is controlled by its induction from activated APCs via Signals 2 and 3 and then maintained by a series of feedback loops. This control of c-Myc is modeled in STORE.2 such that overall quantity of c-Myc per cell is quantified as *M*_*cMYC*_ (Fig. 3). The ability of activated APCs to provide Signals 2 and 3 to T cells over time has been experimentally observed to follow a skewed pulse (*47, 48*), which is approximated using a lognormal distribution with a mean of *μ*_*cyt*_ and standard division of *σ*_*cyt*_:

**Figure 3.**
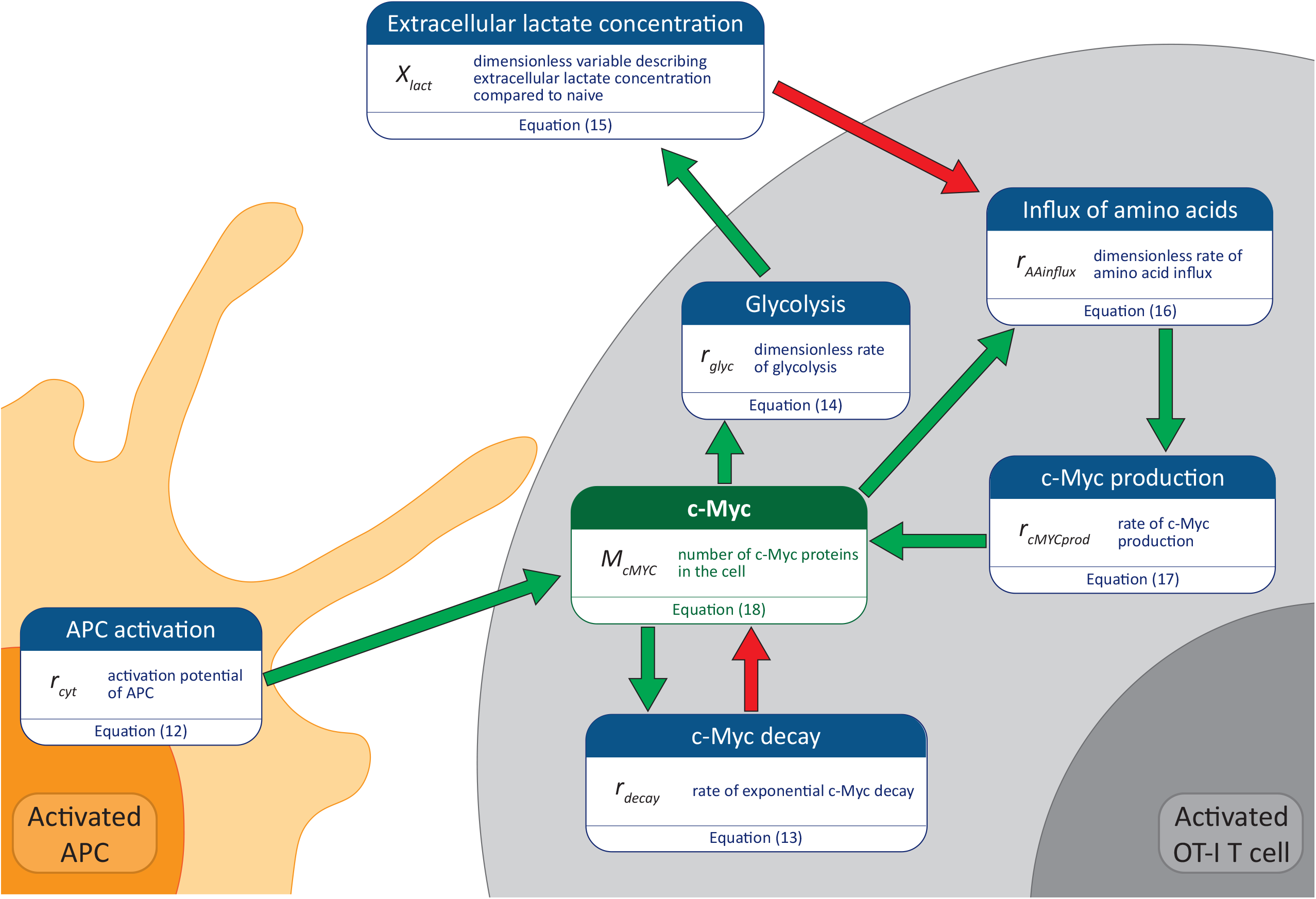
STORE.2 feedback loops controlling c-Myc protein. The protein level of c-Myc in individual CD8^+^ T cells is controlled by a series of feedback loops. While bound to an APC, T cells receive Signals 2 and 3 to enhance c-Myc expression via APC activation. c-Myc upregulation increases the rate of amino acid uptake to increase protein synthesis and c-Myc production. c-Myc also drives increased glycolysis that results in the release of extracellular lactate, which impairs protein synthesis as modeled by the impairment of amino acid influx. The proteolytic degradation of c-Myc is captured by exponential c-Myc decay.

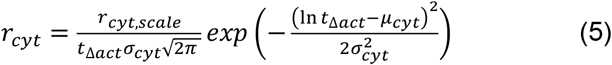

where *t*_Δ*act*_ is the number of hours that have passed since APC activation and *r*_*cyt*_ is the rate of stimulation by an active APC with units of c-Myc molecules per timestep. Importantly, the activation level of APCs is captured as the model parameter *r*_*cyt,scale*_, which is estimated via an optimization-based parameter estimation algorithm for each experimental condition. To translate APC activation to T cell c-Myc synthesis, equation 5 is applied only when T cells are bound to APCs (*AP* and *BP* cell types).

After APC stimulation, the amount of c-Myc present in T cells is determined by its rate of synthesis versus its decay rate. The decay rate of c-Myc in all activated T cells (cells *AP, BP*, and *C* through *J*) is controlled by post-translational modifications that mark the protein for degradation (*35*) as captured in the following formula:

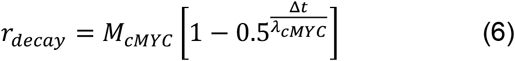

where *λ*^*cMYC*^ is the c-Myc half-life and *r*^*decay*^ is the rate of c-Myc molecules decayed per hour using *M*^*cMYC*^ from the beginning of the timestep and Δ*t* is the length of the timestep.

The process of T cell activation and expansion requires rapid energy production provided by the c-Myc-dependent metabolic switch to glycolysis (*27*) and upregulation of amino acid transporter proteins to enhance uptake of amino acids for protein synthesis (*29*). While glycolysis has the benefit of rapidly producing the energy required for cell growth and division, it has the secondary effect of generating lactate via lactic acid that is secreted into the extracellular fluid. The glycolytic rate (*r*_*glyc*_) and lactate production of each T cell was modeled as follows:

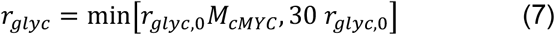

where *r*_*glyc*_ has units of dimensionless lactate concentration production per hour and *r*_*glyc*,0_ is an estimated model parameter that translates the T cell c-Myc level to its glycolytic rate. The upper bound of *r*_*glyc*_ is constrained to be 30 times *r*_*glyc*,0_. The relative lactate concentration (dimensionless) of the extracellular space is then modeled as:

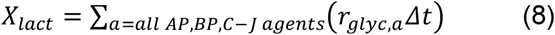

where *X*_*lact*_ is a dimensionless number that represents the extracellular lactate concentration, *r*_*glyc,a*_ is the glycolytic rate for each individual agent indexed by *a. X*_*lact*_ is computed each timestep without any influence by the previous timestep to model the natural, rapid buffering and flow of interstitial fluid in tissues.

The rate of protein synthesis in activated T cells is dependent on the amount of c-Myc in the T cell via upregulation of amino acid transporters (*29*) as well as the amount of lactate in the extracellular fluid (*40, 43*) and is modeled as:

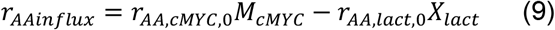

where *r*_*AA,cMYC*,0_ is the rate of amino acid influx per timestep facilitated per c-Myc protein in the cell and *r*_*AA,lact*,0_ is the amount protein synthesis is hindered per dimensionless lactate concentration of the extracellular space. Both parameters were estimated with the parameter estimation algorithm. The synthesis of c-Myc protein from amino acid uptake for each cell was then calculated:

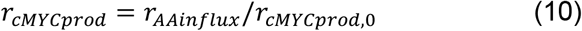

where *r*_*cMYCprod*,0_ is the number of amino acids per c-Myc protein (*r*_*cMYCprod*,0_ *=* 439). It is assumed that c-Myc is not produced during mitosis, therefore *r*_*cMYCprod*_ = 0 during the mitosis (M) phase for all cell types *BP* and *C* through *J*.

Finally, the stimulation by APCs and maintenance of c-Myc through feedback loops are combined to calculate the amount of c-Myc in each AP, BP, and C through J cell type at each time step:

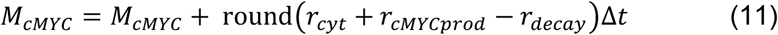

### Modeling the c-Myc levels after initial T cell activation at 24 hr

The induction of c-Myc in T cells has been shown to be increased when the stimulatory capacity of APCs is enhanced. To test the ability of the model to recapitulate the amount of c-Myc protein per OT-I T cell with varying amounts of stimulus, the expression of c-Myc in OT-I/Myc^GFP/GFP^ T cells was measured at 24 hr after activation with peptide alone or in combination with Signal 2 and Signal 3 (Fig. 4A). Measurement of Myc^GFP^ fluorescence showed that the addition of αCD28 (Signal 2) to stimulation with N4 peptide enhanced c-Myc expression while the addition of recombinant IL-12p70 (IL-12, Signal 3) had no additional effect (Fig. 4B, C). To model the c-Myc protein level at 24 hr, the value for *r*_*AA,cMYC*,0_ *in vitro* was estimated by the parameter estimation algorithm to be slightly decreased compared to *in vivo* settings (Table 1). The model was able to capture these changes in c-Myc protein expression by increasing the value of *r*_*cyt,scale*_ with increasing levels of stimulation: N4 peptide = 1, N4 + aCD28 = 35.1, and N4 + aCD28 + IL-12 = 35.3 (Fig. 4D). Importantly, the number of c-Myc proteins per T cell simulated for activation with N4 + aCD28 approximated the copy number of c-Myc per T cell measured by mass spectrometry (*29*), demonstrating that STORE.2 results are physiologically relevant.

**Figure 4.**
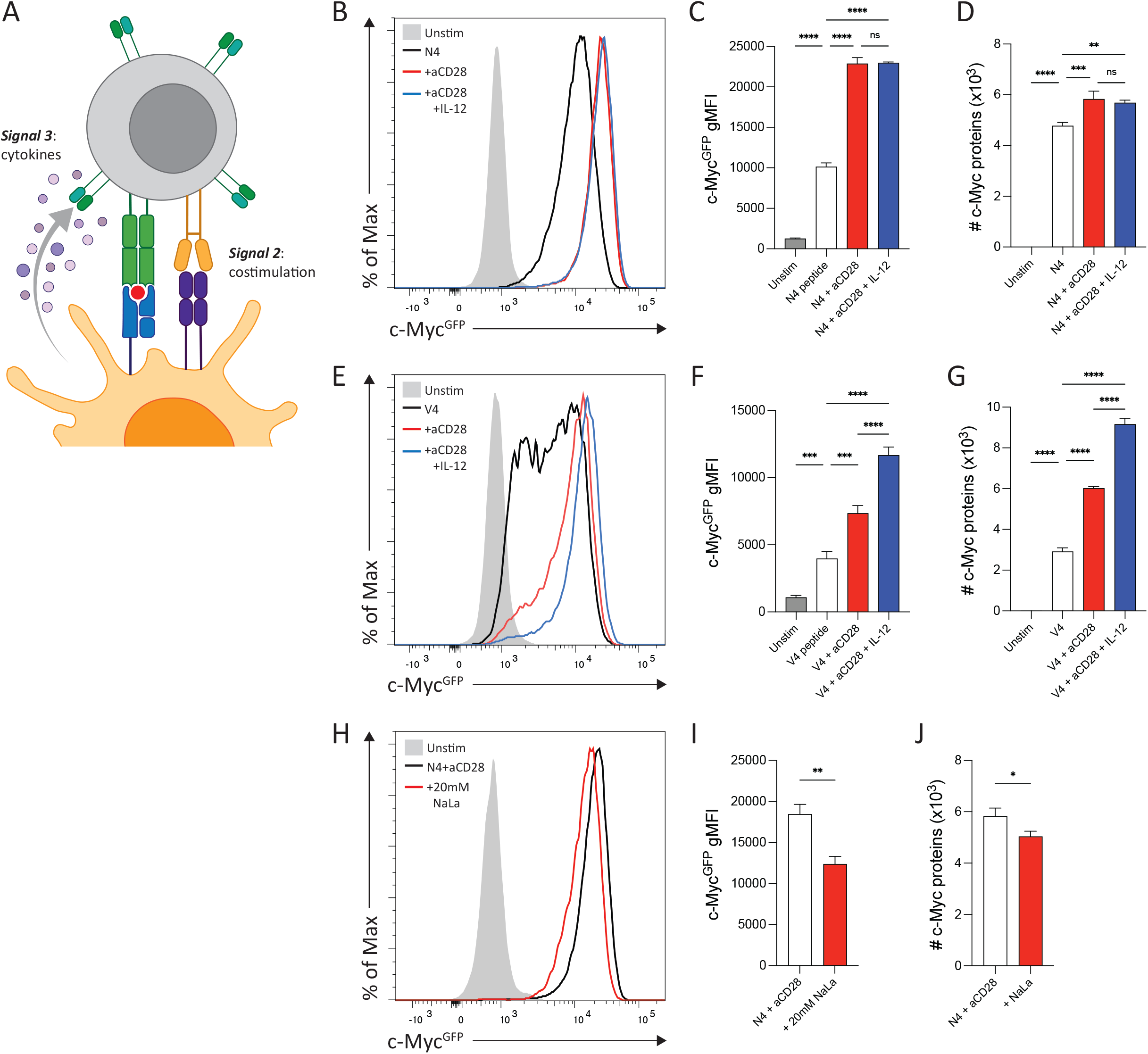
Impact of Signals 2 and 3 on c-Myc expression. (**A**) Schematic of costimulation (Signal 2) and cytokines (Signal 3) acting on a CD8^+^ T cell. (**B**) Flow cytometry analysis of c-Myc^GFP^ fluorescence intensity in OT-I/Myc^GFP/GFP^ T cells stimulated with 20 ng N4 peptide alone, N4 + 2 *μ*g aCD28, or N4 + aCD28 + 1 ng recombinant IL-12p70 (IL-12) *in vitro* for 24 hr compared to unstimulated controls. (**C**) gMFI of c-Myc^GFP^ expression for each stimulation condition with N4 peptide. (**D**) STORE.2 simulation results of c-Myc protein number per OT-I T cell stimulated with N4 peptide, N4 + aCD28, or N4 + aCD28 + IL-12. Parameter estimation resulted in changes to *r*_*cyt,scale*_ for each condition: N4 peptide = 1, N4 + aCD28 = 35.1, and N4 + aCD28 + IL-12 = 35.3. (**E**) Flow cytometry analysis of c-Myc^GFP^ fluorescence intensity in OT-I/Myc^GFP/GFP^ T cells stimulated with 20 ng V4 peptide alone, V4 + 2 *μ*g aCD28, or V4 + aCD28 + 1 ng IL-12 *in vitro* for 24 hr compared to unstimulated controls. (**F**) gMFI of c-Myc^GFP^ expression for each stimulation condition with V4 peptide. (**G**) STORE.2 simulation results of c-Myc protein number per OT-I T cell stimulated with V4 peptide, V4 + aCD28, or V4 + aCD28 + IL-12. Parameter estimation resulted in changes to *r*_*cyt,scale*_ for each condition: V4 peptide = 1, V4 + aCD28 = 175, and V4 + aCD28 + IL-12 = 491. (**H**) Flow cytometry analysis of c-Myc^GFP^ fluorescence intensity in OT-I/Myc^GFP/GFP^ T cells stimulated with N4 + aCD28 +/-20 mM NaLa *in vitro* for 24 hr compared to unstimulated controls. (**I**) gMFI of c-Myc^GFP^ expression for both conditions. (**J**) STORE.2 simulation results of c-Myc protein number per OT-I T cell stimulated with N4 + aCD28 +/-20 mM sodium lactate (NaLa). Parameter estimation showed that increasing *X*_*add*_ to 2.5 x10^9^ recapitulated the effect of 20 mM NaLa. Data are representative plots from (B, C, E, F, H, I) two experiments or (D, G, J) three simulation runs. Bar graphs indicate means ± SD. Data analyzed by (C, D, F, G) one-way ANOVA or (I, J) unpaired *t* test. ns = not significant, \**P* < 0.05, \*\**P* < 0.01, \*\*\**P* < 0.001, \*\*\*\**P* < 0.0001.

In contrast with N4 peptide stimulations, both the addition of Signal 2 and Signal 3 to OT-I/Myc^GFP/GFP^ T cells stimulated with the low affinity V4 peptide increased the expression of Myc^GFP^ (Fig. 4E, F). Despite a difference in the initial step change increase in c-Myc induced by V4 compared to N4 peptide, the model was able to capture the increase in c-Myc expression by only varying the value of *r*_*cyt,scale*_: V4 peptide = 0, V4 + aCD28 = 175, V4 + aCD28 + IL-12 = 491 (Fig. 4G).

The control of c-Myc concentration in T cells by feedback loops includes a suppression of c-Myc synthesis by the presence of lactate. To examine the impact of changes in lactate concentration on c-Myc expression, 20 mM sodium lactate (NaLa) was added to OT-I/Myc^GFP/GFP^ T cells stimulated with N4 peptide and aCD28. Quantification of Myc^GFP^ intensity 24 hr after stimulation showed that expression was significantly decreased in the presence of NaLa. The addition of NaLa was modeled by modifying equation (8) to include the dimensionless *X*_*add*_ term that accounts for externally added lactate concentration:

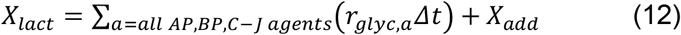

To model the addition of 20 mM NaLa to the culture media, the *X*_*add*_ term was determined to be 2.5 × 10^9^ by parameter estimation. The addition of *X*_*add*_ replicated the impact of NaLa on c-Myc concentration levels in T cells. These data demonstrate that the induction of c-Myc and control of its cellular concentration by multiple feedback loops can be accurately modeled at early stages of T cell activation.

### Cell cycle progression tracking

In contrast with the STORE.1 model, STORE.2 no longer uses PDFs to select the amount of time it takes for an activated T cell to divide. Instead, STORE.2 tracks the progress of each activated T cell through the G1, S, G2, and M phases of cellular mitosis using a new value called *M*_*cycle*_. *M*_*cycle*_ is a number between 0% (which is the beginning of the G1 phase) and 100% (which is the beginning of the M phase). During this early phase of priming and expansion of the T cell response, all T cells are assumed to begin at G1 (0%) immediately upon division. Progression through the cell cycle is then a function of c-Myc levels in the T cell as follows:

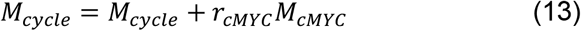

where *r*_*cMYC*_ is a proportionality constant with units of % per c-Myc protein in the cell per timestep. The value of *r*_*cMYC*_ is estimated in the parameter estimation step. After *M*_*cycle*_ for the T cell has reached 100%, the cell enters the M phase, which is approximated to take 1 hr (*49*). Once the M phase is complete, the model assumes that the cells divide evenly in half with each daughter cell getting one half of the c-Myc protein in the parent.

### Modeling c-Myc dependent T cell division

The division destiny of CD8^+^ T cells has been shown to be dependent the degree of stimulation by APCs (*22*–*24*) that drives increasing amounts of c-Myc (*24*). To determine whether the addition of c-Myc induction and maintenance to STORE.2 would allow for the modeling of stimulation-dependent T cell division, purified OT-I/Myc^GFP/GFP^ T cells were labeled with CellTrace Violet (CTV), stimulated with peptide in combination with Signals 2 and 3, and allowed to proliferate for 72 hr *in vitro* (Fig. 5A). Stimulation with N4 peptide alone resulted in robust T cell division with a mean division number (MDN) of 3.00±0.05. The addition of Signal 2 increased this MDN to 3.78±0.04, while the addition of Signal 3 had no further impact on division (MDN = 3.53±0.03) (Fig. 5B). These differences in MDN were accurately predicted by the difference in c-Myc^GFP^ expression at 24 hr (Fig. 4C). To visualize division number with the model, each *A* cell is assigned a “cell dye” value of 4096 and cells in subsequent divisions receive a cell dye value of 4096/2^*n*^ where *n* is the division number. Simulations of the T cell response at 72 hr showed that the model could match what was measured experimentally (Fig. 5C) as the c-Myc levels at 24 hr also predicted the differences in the MDN at 72 hr (Fig. 5D). The changes to *r*_*cyt,scale*_ that drove the c-Myc level differences at 24 hr also controlled differences in MDN at 72 hr, similar to experimental changes in stimulation level. Experimentally, stimulation with V4 peptide alone resulted in fewer divisions (MDN = 1.97±0.09) than stimulation with N4 peptide (Fig. 5E). Addition of Signal 2 to V4 stimulation resulted in an increased MDN (3.40±0.06) compared to peptide alone, and the addition of Signal 3 further increased MDN (4.11±0.09) (Fig. 5F). Similar to the N4 peptide, modeling of V4 peptide in combination with Signals 2 and 3 was able to capture this dynamic with changes in *r*_*cyt,scale*_ alone (Fig. 5G-H).

**Figure 5.**
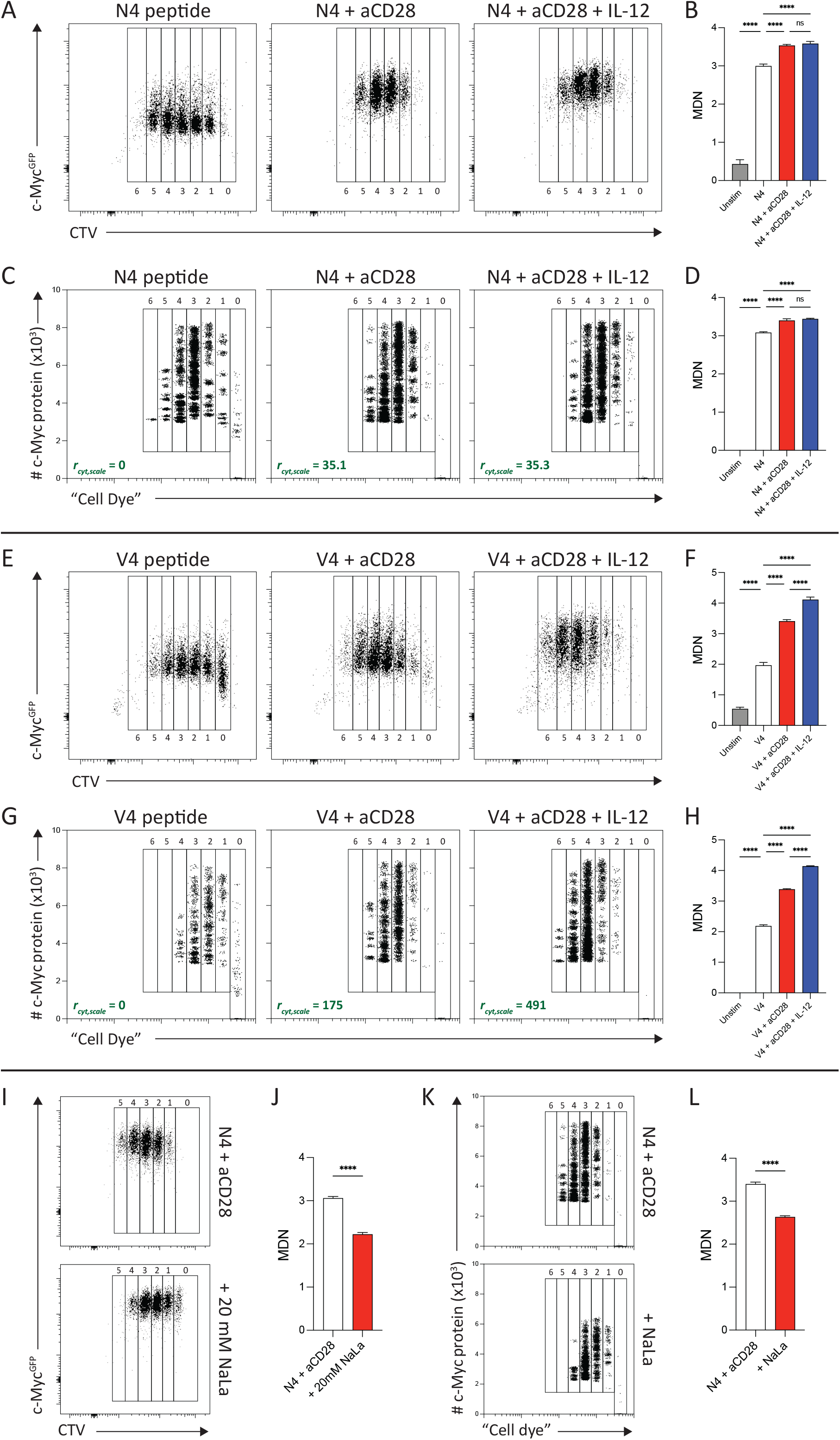
Role of T cell stimulation in T cell division. (**A**) Flow cytometry analysis of CTV-labeled OT-I/Myc^GFP/GFP^ T cells stimulated with 20 ng N4 peptide alone, N4 + 2 *μ*g aCD28, or N4 + aCD28 + 1 ng IL-12 *in vitro* for 72 hr. (**B**) Mean division number (MDN) for each N4 stimulation condition at 72 hr. (**C**) STORE.2 simulation results of c-Myc protein number and T cell division measured by dilution of “cell dye” at 72 hr. Insets show value of *r*_*cyt,scale*_ for each condition. (**D**) MDN calculated from STORE.2 simulations of different N4 stimulation conditions. (**E**) Flow cytometry analysis of CTV-labeled OT-I/Myc^GFP/GFP^ T cells stimulated with 20 ng V4 peptide alone, V4 + 2 *μ*g aCD28, or V4 + aCD28 + 1 ng IL-12 *in vitro* for 72 hr. (**F**) MDN calculated for each V4 stimulation condition at 72 hr. (**G**) STORE.2 simulation results of c-Myc protein number and T cell division at 72 hr. Insets show value of *r*_*cyt,scale*_ for each condition. (**H**) MDN calculated from STORE.2 simulations of different V4 stimulation conditions. (**I**) Flow cytometry analysis of CTV-labeled OT-I/Myc^GFP/GFP^ T cells stimulated with N4 + aCD28 +/-20 mM NaLa *in vitro* for 72 hr. (**J**) MDN calculated for N4 + aCD28 +/-20 mM NaLa. (**K**) STORE.2 simulation results of c-Myc protein number and T cell at 72 hr for OT-I T cell stimulated with N4 + aCD28 +/-NaLa (*X*_*add*_ = 2.5 × 10^9^). (**L**) MDN calculated for N4 + aCD28 +/-NaLa. Data are representative plots from (A, B, E, F, I, J) two experiments or (C, D, G, H, K, L) three simulation runs. Bar graphs indicate means ± SD. Data analyzed by (B, D, F, H) one way ANOVA or (J, L) unpaired T-test. ns = not significant, \*\*\*\**P* < 0.0001.

To test the impact of the lactate negative feedback loop of c-Myc synthesis on T cell division, OT-I/Myc^GFP/GFP^ T cells were stimulated with N4 peptide and Signal 2 in the presence of 20 mM NaLa. Analysis of T cell division at 72 hr after stimulation showed that T cells proliferated less in the presence of NaLa (Fig. 6I) with a significant decrease in MDN (2.22±0.04) compared to N4 + aCD28 alone (3.06±0.04) (Fig. 5J). To model these conditions, *X*^*add*^ was increased from 0 to 2.5 × 10^9^ to account for the addition of NaLa to the culture media. Again, this change to a single variable was able to capture the decrease in cell division (Fig. 5K) and MDN (Fig. 5L) observed experimentally. Importantly, these model results are a key validation of the model because these *in vitro* peptide stimulation experiments are extrapolative scenarios outside of the *in vivo* experimental data used for model training in the parameter estimation step. The ability to recapitulate the increased APC stimulatory capacity and addition of lactate with changes to a single first-principles based parameter (*r*_*cyt,scale*_ and *X*_*add*_, respectively) provides evidence of the correctness of the underlying phenomenological structure of STORE.2.

**Figure 6.**
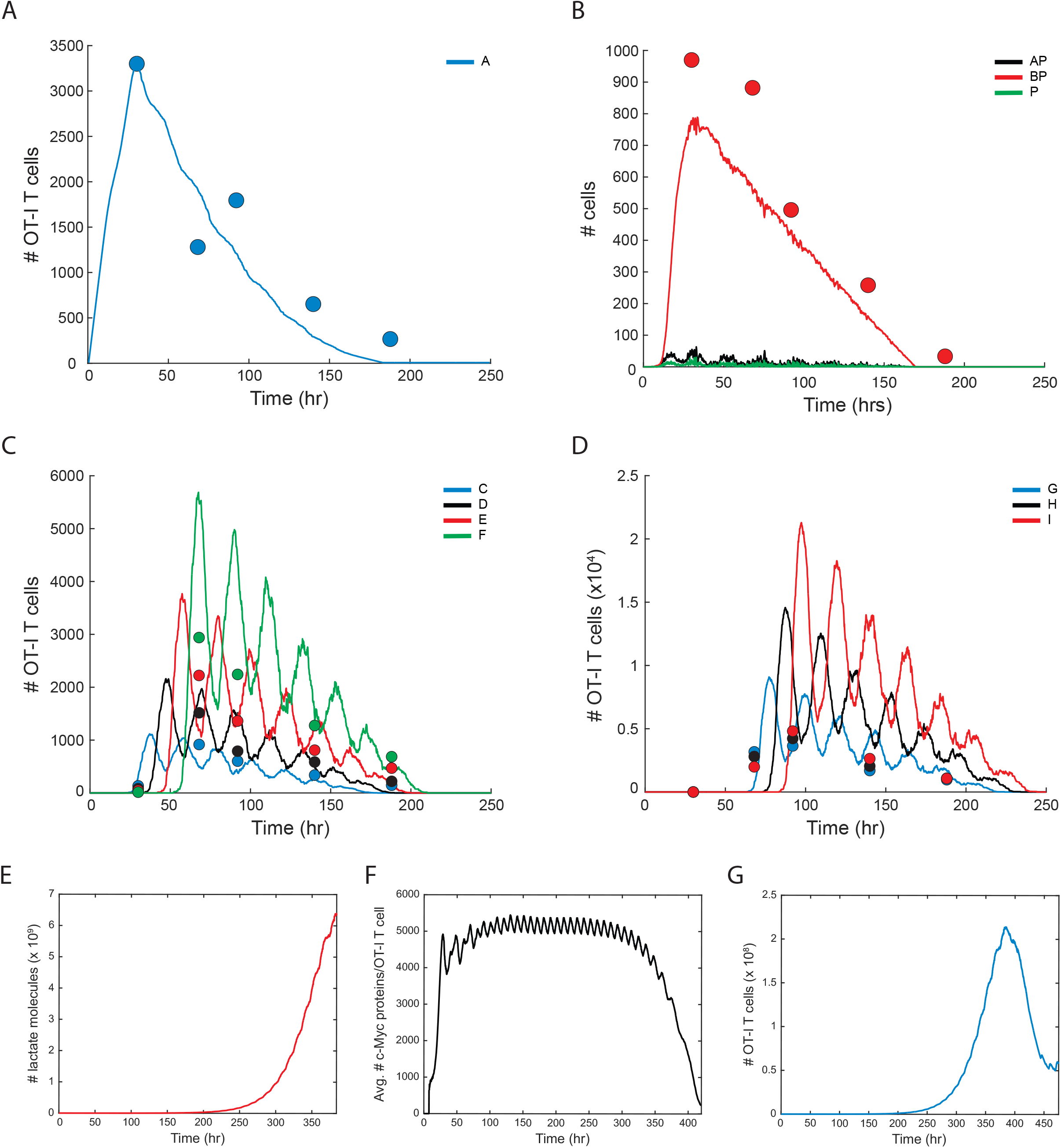
Simulation of the CD8^+^ T cell response to vaccination. Representative plots of a single STORE.2 simulation of the OT-I T cell response to i.p. immunization with CPS-OVA parasites for 250 hours post-immunization. Lines represent simulation results and filled circles represent experimental OT-I cell counts in the omentum measured by flow cytometry. (**A**) Number of naive OT-I T cells (*A*) versus time. (**B**) Number of unbound activated APCs (*P*), bound pairs of *P* and Nur77^GFP-^ OT-I T cells (*AP*), and bound pairs of *P* and Nur77^GFP+^ OT-I T cells (*BP*) versus time. (**C**) Number of OT-I T cells in divisions 1-4 (divisions *C*-*F*) versus time (**D**) Number of OT-I T cells in divisions 5-7 (divisions *G*-*I*) versus time. Data are representative plots from (filled circles) two experiments or (lines) three simulations runs. (**E**) Number of lactate molecules (dimensionless) in the extracellular space versus time. (**F**) Average number of c-Myc proteins per OT-I T cell versus time. (**G**) Number of OT-I T cells versus time. Data are representative plots from (A, B, C, D) two experiments or (A to G) three simulation runs. Filled circles indicate means.

### Modeling T cell responses in vivo

While most computational models examining the early activation phase of the CD8^+^ T cell response are derived from *in vitro* data, the STORE models were designed to simulate the *in vivo* response to vaccination. Thus, STORE.2 was used to model the kinetics of the OT-I T cell response in the omentum of wild-type (WT) mice after intraperitoneal vaccination with the CPS parasite (*38*). The final values of the STORE.2 model parameters were determined by the parameter estimation algorithm and are shown in Table 1. All parameters except one are well inside their bounds, showing that no individual parameter is limiting the performance of the model. The one parameter at a bound (*r*_*glyc,o*_) has only a small impact on the model, and lowering it below the bound would have only a small effect on the model. In addition to the parameter changes to model the *in vitro* priming experiments mentioned above, one key parameter change for the *in vivo* modeling was the decrease of the initial change in c-Myc after Signal 1 (*μ*_*cMYC*0_ = 800, *σ*_*cMYC*0_ = 200). This change could result from processing and presentation of parasite-derived OVA protein by professional APCs *in vivo* rather than self-presentation of N4 peptide by T cells. Example results from a single simulation of the STORE.2 model demonstrate that the model fits the data from the experimental time points for each cell type (Figure 6A-D). The stochasticity of the STORE.2 model accounts for variable behavior of individual cells, but the trends of the cell populations as a whole are consistent between simulations. A key characteristic of the simulation is the “ringing” (oscillating) cell counts of the various agents *C*-*I* once created. For example, cell type *F* goes through at least 6 distinct maxima during the response, roughly 20-24 hours apart. This ringing occurs because the availability of activated APCs (*P*) is limited compared to the available naive T cells (*A*). As each *A* + *P* → *AP* → *BP* → *P* + 2*C* transition sequence is completed (which takes approximately 20-22 hours), the *P* is free to bind another unbound *A* after a short time, starting the next wave. The fact that the c-Myc models of STORE.2 capture this cyclic behaviour in a way that naturally arises from the fundamental phenomena so well is a key finding.

The gradual decay in c-Myc levels per T cell has been implicated in a mechanism for the cessation of T cell division (*24*). While the STORE.2 model was designed to model the expansion phase of the CD8^+^ T cell response, extension to approximately 400 hours post immunization shows the impact of clonal expansion on T cell behavior. The exponential growth in the number of T cells generated during the response results in an equivalent increase in the amount of lactate generated per timestep (Fig. 6E). The buildup of lactate arising from large cell populations causes average c-Myc protein count per cell to decrease (Fig. 6F). Cells with lower c-Myc protein counts still progress through the cell cycle, but lower c-Myc levels at the time of mitosis result in the inheritance of lower starting c-Myc protein counts by both daughter cells. The heritable level of c-Myc protein provides a mechanism for the rapid and population-level slowing of division rates such that the natural leaving rate of later division T cells from the system (*p*_*IJ,leave*_) eventually outpaces the growth rate of the population. Consequently, the number of T cells in the system begins to exponentially decay (Fig. 6G). While the model was not explicitly designed to predict contraction and no data from this period was used in parameter estimation, it can be concluded that the transition from expansion to contraction phases simulated here is at least partially explained by phenomena considered in STORE.2.

## DISCUSSION

Computational models have served important roles in characterizing the dynamics of CD8^+^ T cell responses that are difficult to measure experimentally (*11*). More recently, these models have been adapted to identify and track the role of heritable factors that impact the expansion of specific clones in CD8^+^ T cell responses. Thus, models have moved from describing population-based responses to tracking specific clones using ABM (*26*). STORE.2 is a stochastic, ABM that captures the differences in the TCR signal strength that are critical for the expansion rate and DD of CD8^+^ T cells by modeling the induction of the transcription factor c-Myc after Signal 1. STORE.2 also uniquely incorporates the impact of the T cell extrinsic factors including Signals 2 and 3 from APCs and the extracellular environment by modeling their role in c-Myc expression levels in each T cell. Importantly, STORE.2 uses this c-Myc expression rather than probability distributions as a heritable factor that determines the division time and DD of each T cell. By combining the ability to track the roles of Signals 1-3 on c-Myc expression with agent-based modeling of APCs, future work with STORE.2 can explicitly track the roles of peptide affinity and APC stimulatory capacity in the development of clonal T cell responses.

The STORE.1 model introduced the first ABM to describe the behavior of both the T cell and APC. This APC tracking allowed STORE.1 to capture the timing of antigen presentation via the arrival time of *P* into the system and the exit rate of *BP* out of the system (*BP*_*leave*_). STORE.2 expanded this model by adding the time required for APCs to become activated and capable of presenting cognate antigen, as well as the stimulatory capacity of these activated APCs when they are bound to T cells as either AP or BP cell types (*r*_*cyt,scale*_). These parameters describing APC behavior now allow for the model to simulate the kinetics of different vaccine formulations. For example, the rapidity and duration of APC antigen presentation is a critical factor for vaccine efficacy and is variable between vaccine formulations. For example, subunit vaccines provide short-lived, single dose of antigen while mRNA vaccines and vaccines that use attenuated microbes utilize mechanisms that increase both the load and duration of antigen available to activate CD8^+^ T cells. Another critical element in developing T cell vaccines is the ability of adjuvants to engage different stimulatory pathways that support robust CD8^+^ T cell responses. The addition of *r*^*cyt,scale*^ and the use of c-Myc as a readout for T cell stimulus provides a means to quantify the stimulatory capacity of T cell vaccines and predict the resulting CD8^+^ T cell response. Future work will use this agent-based APC modality to model the efficacy of different adjuvant combinations in generating protective CD8^+^ T cell vaccines.

The STORE.2 model uniquely models the interplay between the T cells in the priming site and the extracellular inflammatory conditions. As T cells go through the metabolic switch to glycolysis in the site of priming and begin secreting lactic acid, previous work has shown that draining LN becomes acidic (*44*). This lactic acid results in an increase in lactate concentrations that are capable of impairing T cell metabolism and proliferation (*41*–*44*), which is captured in the model by controlling the amount of c-Myc in the cell through changes in the rate of c-Myc synthesis. This unique functionality of STORE.2 could have important applications in understanding differences in T cell responses generated in environments of high cellular stress. For example, lactate concentrations in solid tumors are significantly higher than in healthy secondary lymphoid organs, which has been shown to impair T cell responses (*50*). The STORE.2 model would be able to account for this suppressive environment and provide predictive insights into designing better anti-cancer T cell therapies. In summary, STORE.2 succeeds in replacing simple assumed behaviors for cell division in STORE.1 with a first-principles, dynamic system model that explains the fundamental phenomena of c-Myc-dependent T cell activation and proliferation. STORE.2 thus mechanistically captures the role of T cell extrinsic factors in generating T cell responses to provide an accurate and predictive model of CD8^+^ T cell activation after vaccination.

## METHODS

### Model Implementation

The STORE.2 model was implemented in MATLAB. The run time of a single simulation takes typically about 30 seconds.

### Parameter Estimation

In the prior work, the STORE.1 model parameter values were estimated through optimization techniques that compared against *in vivo* experimental data (*38*). In this work, the parameters new to STORE.2 were estimated while leaving the parameters common to STORE.1 at their values determined in the prior work. The same experimental data set used in the prior work was used as the training set in the present work. An optimization procedure was used to estimate 12 of the 14 parameters new to the STORE.2 model (two parameters were fixed based on literature values, see Table 1). The associated optimization problem is to determine the new STORE.2 model parameters that minimizes of the relative error between the cell counts of various types of agents and at various time points predicted by the STORE.2 model versus the *in vivo* experimental data. The decision variables were the 12 unknown model parameters. The equality constraints are the black-box STORE.2 model as executed by the MATLAB implementation. The objective function (the relative error between simulation and experiment) is defined as:

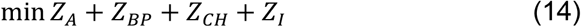

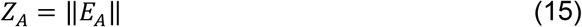

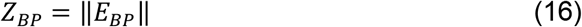

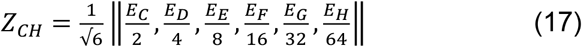

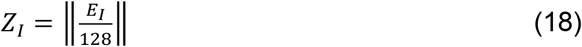

where the four *Z* variables are scalars, and are the normalized, weighted error between simulation results and experimental data points for four groups of cell types: A, BP, C through H, and I, such that each of the four *Z* groups are weighted equally in the final objective function metric. Cell type J was not considered in the objective function and therefore not used for training the parameters; it is left as a testing metric instead.

The error *E*_*i*_ of cell type *i* is:

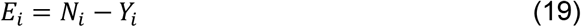

where *N*_*i*_ is the vector of cell counts of cell type *i* at each simulation time point, and *Y*_*i*_ is the corresponding experimentally determined cell counts at those same time points. The optimization problem was solved using the wild-type (WT) residing mice data from *in vivo* experiments in (*38*) as the *Y*_*i*_.

The particle swarm optimization (PSO) algorithm built into MATLAB was used as the optimization algorithm, with 50 particles and 100 iterations. It was executed on a Dell XPS 13 (790) computer with a 10^th^ generation Intel processor running parallelized with 4 workers. PSO cannot guarantee global minimization, but the procedure was repeated several times to reduce the chance of missing the region of the global minima. Each PSO run required approximately 3-6 hours (wall time) to complete.

### Mice

All procedures involving mice were reviewed and approved by the Institutional Animal Care and Use Committee of the University of Pennsylvania (Animal Welfare Assurance reference number A3079-01) and were in accordance with the guidelines set forth in the Guide for the Care and Use of Laboratory Animals of the National Institutes of Health (NIH).

C57BL/6J (stock no. 000664, RRID: IMSR_JAX:000664), Nur77^GFP^ (stock no. 016617, RRID:IMSR_JAX:016617), OT-I (stock no. 003831, RRID:IMSR_JAX:003831), Myc^GFP^ (stock no. 019075, RRID: IMSR_JAX:019075) mice were obtained from the Jackson Laboratory.

### Immunizations and infections

All *T. gondii* immunizations were performed using 2 × 10^5^ CPS-OVA parasites 2 hours after OT-I T cell transfer as previously described (*38*). CPS-OVA parasites have been previously described (*51*) and were derived from the RHΔ*cpsII* clone, which was provided as a gift by D. Bzik. Parasites were cultured and maintained by serial passage on human foreskin fibroblast cells in the presence of parasite culture media [71.7% Dulbecco’s Modified Eagle’s Medium (Corning, 10-017-CM), 17.9% Medium 199 (Gibco, 11150-059), 9.9% fetal bovine serum (FBS) (Atlanta Biologicals, S11150H), 0.45% penicillin and streptomycin (final concentration of penicillin: 0.05 U/ml and streptomycin: 50 μg/ml; Gibco, 15140-122), and 0.04% gentamicin (final concentration of gentamicin: 0.02 mg; Gibco, 15750-060)], which was supplemented with uracil (final concentration of 0.2 mM uracil; Sigma-Aldrich, U1128). For infections, parasites were harvested and serially passed through 18-and 26-gauge needles (BD, 305196 and 305115) before filtration with a 5-μm filter (PALL 4650 Acrodisc). Parasites were washed extensively with PBS, and mice were injected intraperitoneally with parasites suspended in PBS.

### OT-I T cell enrichment and tissue harvesting

For OT-I T cell enrichment, OT-I mice were interbred with CD45.1/Nur77^GFP^ mice or Myc^GFP/GFP^ mice. To isolate OT-I CD8^+^ T cells, LNs and spleens were harvested, and leukocytes were obtained by processing spleens and LNs over a 70-μm filter (Fisher Scientific, 22-363-548) and washing them in complete RPMI [90% RMPI 1640 (Corning, 10-040-CM), 10% FBS, 1% penicillin-streptomycin, 1 mM sodium pyruvate (Corning, 25-000-Cl), 1% non-essential amino acids (Gibco, 11140-050), and 0.1% β-mercaptoethanol (Gibco, 21985-023)]. Red blood cells were then lysed by incubating for 5 min at room temperature in 5 ml of lysis buffer [0.864% ammonium chloride (Sigma-Aldrich, A0171) diluted in sterile deionized H2O], followed by washing with complete RPMI. OT-I T cells were then purified by magnetic activated cell sorting (MACS) using the CD8a+ T Cell Isolation Kit (Miltenyi Biotec, 130-104-075). Purified OT-I T cells were then fluorescently labeled using the CTV (Thermo Fisher Scientific, C34557) labeling kit. For *in vivo* studies, OT-I T cells were then transferred by intraperitoneal injection into recipient mice.

Peritoneal exudate cells were obtained by peritoneal lavage with 8 ml of ice-cold PBS. Omentum was isolated, incubated in LiberaseTL (0.4 U/ml; Roche, 5401020001) for 1 hour at 37°C, passed through an 18-gauge needle, and processed over a 70-μm filter. Leukocytes from the spleen and DLNs were obtained by processing spleens and LNs, washing them in complete media, and lysing red blood cells (see above). Cells were then resuspended in complete RPMI.

### OT-I T cell activation in vitro

For *in vitro* assays measuring OT-I T cell priming and expansion, MACS-purified, CTV-labeled OT-I/Myc^GFP/GFP^ T cells were plated in a round bottom 96 well plate at 10^5^cells in 200 *μ*L per well. T cells were stimulated with combinations of 20 ng per well of either SIINFEKL (N4) or SIIVFEKL (V4) peptide (Anaspec), 2 *μ*g per well anti-CD28 (BioXcell, BE0015-1), and 1ng per well recombinant murine IL-12p70 (Peprotech, 210.12). The effect of IL-2 made from the OT-I T cells was blocked using 2 *μ*g of anti-IL-2 antibody (Clone S4B6, BioXcell, B0043-1). To test the effect of lactate on c-Myc expression and T cell division, sodium lactate (MilliporeSigma, 71718) was added to each well at 0 hours to a final concentration of 20 mM. Cells were incubated for 24, 48, and 72 hours before analysis by flow cytometry. Mean division number (MDN) was calculated by 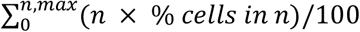, *n* = division number, *n, max* = maximum number of measured divisions.

### Flow cytometry

Cells were washed with FACS buffer [1× PBS, 0.2% bovine serum antigen (Gemini, 700-100P), and 1 mM EDTA (Gibco, 15575-038)] and incubated in Fc block [99.5% FACS buffer, 0.5% normal rat immunoglobulin G (Invitrogen, 10700), and 2.4G2 (1 μg/ml; Bio X Cell, BE0307)] at 4°C for 10 min before staining. If cells were stained for cell death using LIVE/DEAD staining, then LIVE/DEAD Fixable Aqua Dead Cell marker (Invitrogen, L34957), Ghost Dye Violet 510 Viability Dye (Tonbo Biosciences, 13-0870-T100), or Ghost Dye Red 780 Viability Dye (Tonbo Biosciences, 13-0865-T100) was included during incubation with Fc block. Cells were surface-stained in 50 μl at 4°C for 15 to 20 min and washed in FACS buffer before acquisition. For intracellular transcription factor staining, cells were rinsed with FACS buffer and surface-stained as described above, fixed using the eBioscience Foxp3 Transcription Factor Fixation/Permeabilization Concentrate and Diluent (Thermo Fisher Scientific, 00-8222) for 30 min at 4°C, and then washed with FACS buffer. Cells were then stained for intracellular cytokines and transcription factors in 50 μl of 1× eBioscience Permeabilization Buffer (Thermo Fisher Scientific, 00-8333-56) at 4°C for at least 1 hour. Cells were then washed in FACS buffer before acquisition.

The following antibodies were used for staining: CD11a (PerCp-Cy5.5; BioLegend, 101124; clone: M17/4; RRID:AB_2562932), CD8a (BV605; BioLegend, 100744; clone: 53-6.7; RRID:AB_2562609), CXCR3 (BV650; BioLegend, 126531; clone: CXCR3-173; RRID:AB_2563160), KLRG1 (BV711; BioLegend, 138427; clone: 2F1/KLRG1; RRID:AB_2629721), CD3 (BV785; BioLegend, 100232; clone: 17A2; RRID:AB_2562554), CD25 (APC; eBioscience, 17-0251-82; clone: PC61.5; RRID:AB_469366), CD4 (AF700; BioLegend, 100536; clone: RM4-5; RRID:AB_493701), CD45.1 (APC-ef780; Invitrogen, 47-0453-82; clone: A20; RRID:AB_1582228), TCR Vα2 (PE; BioLegend, 127808; clone: B20.1; RRID:AB_1134183), CD69 (PE-Cy7; BD Biosciences, 552879; clone: H1.2F3; RRID:AB_394508), aGFP (AF488; BioLegend, 338008; clone: FM264G; RRID:AB_2563288), CD8a (BUV395; BD Biosciences, 563786; clone: 53-6.7; RRID:AB_2732919), CD11a (BUV805; BD Biosciences, 741919; clone: 2D7; RRID:AB_2871232), CD25 (BV785; BioLegend, 102051; clone: PC61; RRID:AB_25641311), CD62L (BV711; BioLegend, 104445; clone: MEL-14; RRID:AB_2564215), CD69 (BUV737; BD Biosciences, 612793; clone: H1.2F3; RRID:AB_2870120), CD71 (PE-Dazzle594; BioLegend, 113817; clone: RI7217; RRID:AB_2749883), CD98 (PE-Cy7; BioLegend, 128214; clone: RL388; RRID:AB_2750547), CD122 (PE-Cy5; BioLegend, 123220; clone: TM-B1; RRID:AB_2715962), cMyc (PE; Novus, NB600-302; clone: 9.00E+10), Ki67 (RB613; BD Biosciences, 571126; clone: B56; RRID:AB_3686255), O-Glcnac (AF647; ThermoFisher Scientific, 51-9793-42; clone: RL2; RRID:AB_2784743). Samples were run on a BD FACSymphony A3 (BD) and analyzed using FlowJo Software (Tree Star).

## STATISTICAL ANALYSIS

Statistical analysis was performed using GraphPad Prism 10 software. The statistical analysis and significance values for each comparison are indicated in each figure caption.

## ACKNOWLEDGEMENTS

The flow cytometry data for this manuscript were generated in the Penn Cytomics and Cell Sorting Shared Resource Laboratory at the University of Pennsylvania (RRID:SCR_022376). Penn Cytomics is partially supported by the Abramson Cancer Center NCI Grant (P30 016520). This work was supported by a grant from the National Institutes of Health: NIAID AI160664.

